# The consequences of host heterogeneity for parasite transmission depend on how host infectiousness is determined

**DOI:** 10.1101/2024.09.16.613211

**Authors:** Jacob A. Cohen, Andrew D. Dean, Mark Viney, Andy Fenton

**Affiliations:** Department of Biology, University of Oxford, 11a Mansfield Road, Oxford OX1 3SZ, UK; Department of Evolution, Ecology and Behaviour, Institute of Infection, Veterinary and Ecological Sciences, University of Liverpool, Liverpool L69 7ZB, UK

**Keywords:** disease ecology, host heterogeneity, parasite transmission, susceptibility, infectiousness, epidemiological model

## Abstract

Host heterogeneities in susceptibility and infectiousness can affect parasite transmission, but standard analyses typically consider a limited range of epidemiological scenarios. We propose two contrasting phenomenological scenarios that capture a wide range of processes by which host infectiousness is determined, and explore how these scenarios affect the impact of host heterogeneities on parasite transmission. Specifically, we contrast ‘Recipient dependence’ (RD), where a host’s infectiousness is a fixed characteristic of the recipient host being infected, with ‘Donor dependence’ (DD), where a host’s infectiousness is determined by the donor host that infected them. Contrasting Susceptible-Infected models of these two phenomenological scenarios show that under RD, *R*_*0*_ is driven by population-level covariance between susceptibility and infectiousness, but for DD scenarios it is driven by the maximum infectiousness present in the population. Our results show that these different scenarios, which capture different ways by which host infectiousness is determined, should be considered explicitly when exploring the consequences of host heterogeneity on transmission.

## Introduction

Individuals can vary substantially in their propensities to be infected by and transmit parasites (VanderWaal *et al*. 2016). These individual-level host heterogeneities can have significant effects on the transmission of parasites and so affect patterns of infection in host populations. Superspreaders – hosts that are responsible for a disproportionate number of transmission events – are a well-known, if extreme, example of such host heterogeneity (Woolhouse *et al*. 1997; Lloyd-Smith *et al*. 2005; Lemieux *et al*. 2021). Despite the commonly accepted importance of host heterogeneity, there has been somewhat limited investigation of this, particularly in understanding the epidemiological consequences of how a host’s infectiousness is determined.

Parasite transmission is a function of three factors: (i) host susceptibility, which is the host’s propensity to become infected following parasite exposure, (ii) host infectiousness, the capacity of an infected host to transmit parasites, and (iii) host contact rate, the rate of transmission-relevant contacts, dependent on the transmission mode of the parasite in question (McCallum *et al*. 2017). Variation among individual hosts in any of these traits can alter parasite transmission dynamics in a host population (Dwyer *et al*. 1997; Coutinho *et al*. 1999; Barlow 2000; Matthews *et al*. 2006; Streicker *et al*. 2013; Kong *et al*. 2016; Stephenson *et al*. 2017; White *et al*. 2018). Importantly, heterogeneities in these three traits can exist simultaneously (so-called ‘coupled heterogeneities’), and potentially interact with each other, altering transmission dynamics (Yates *et al*. 2006; Miller 2007; Hickson *et al*. 2014; Vazquez-Prokopec *et al*. 2016). For instance, it has been shown that covariation between host heterogeneities can increase or decrease the parasite’s basic reproduction number, *R*_*0*_, depending on the traits considered and whether the covariation is positive or negative (Vazquez-Prokopec *et al*. 2016; Lloyd *et al*. 2020).

To date, most studies investigating the consequences of host heterogeneities on parasite transmission dynamics implicitly assume that a host’s infectiousness is a fixed characteristic of the ‘recipient’ individual being infected (Coutinho *et al*. (1999); Yates *et al*. (2006); Miller (2007); Lloyd *et al*. (2020)). We call this ‘Recipient dependence’ (RD) because the infectiousness of the newly infected host (the recipient) is solely determined by characteristics of that individual. There are a range of processes which may lead to RD determination of host infectiousness. For example, hosts might differ in terms of genotype (Anacleto *et al*. 2019), nutritional status (Cornet *et al*. 2014), behavioural phenotype (Araujo *et al*. 2016), or social connectedness (Mantzaris *et al*. 2013; Adelman *et al*. 2015), all of which result in levels of infectiousness linked to those characteristics of the recipient host. Generally, most studies of host heterogeneities implicitly assume RD-based scenarios (*e*.*g*. Coutinho *et al*. (1999); Yates *et al*. (2006); Miller (2007); Lloyd *et al*. (2020)). In such systems, heterogeneity can either increase or decrease *R*_*0*_, depending on whether the covariance between host susceptibility and infectiousness is positive or negative (Lloyd *et al*. 2020).

We contrast the RD scenario with scenarios where the infectiousness of a newly infected host depends on characteristics of the ‘donor’ host that infected it. We call this ‘Donor dependence’ (DD) because the infectiousness of the newly infected host is determined by the infecting host (the donor). DD may occur, for example, when the parasite load received by a newly infected host determines how infectious that host becomes (Strong *et al*. 2015), such as when highly infectious hosts generate other highly infectious hosts (Beldomenico 2020; Wanelik *et al*. 2023). It can also capture situations where co-circulating parasite strains differ in their infectivity (Brault *et al*. 2004; Dai *et al*. 2013; Medina *et al*. 2018), so that hosts infected with highly infective parasite strains then become highly infectious hosts themselves. Critically, and in contrast to the RD scenario, in the DD scenario a host’s infectiousness is determined solely by the identity of the donor host that infected them. We propose that RD and DD are two epidemiologically distinct phenomenological ways to conceptualise various processes by which a host’s infectiousness is determined, distinguished by the identity of the host (the recipient or the donor) that determines the infectiousness of a newly infected host.

We suggest that the differentiation of the RD and DD scenarios will lead to different patterns of host infectiousness in a population and, crucially, alter the impact of host heterogeneities on parasite transmission. Previous work (e.g., Vazquez-Prokopec *et al*. (2016); Lloyd *et al*. (2020); Wanelik *et al*. (2023)) has implicitly used one or the other of these scenarios but has not systematically explored how they differ in affecting the impact of coupled heterogeneities in susceptibility and infectiousness on parasite transmission. To investigate this, we develop Susceptible-Infected (SI) compartmental models that incorporate heterogeneity in susceptibility and infectiousness under both RD and DD scenarios. We then analyse these models to determine how RD and DD affect the relationships between those host heterogeneities, parasite transmission and host population dynamics. We find that how infectiousness is determined – RD *vs*. DD – can have substantial effects on parasite transmission, particularly affecting the primary drivers of the basic reproduction number (*R*_*0*_).

## Material and methods

### Model Framework

The standard density-dependent SI model in a homogeneous population of size *N* divided into susceptible (*S*) and infected (*I*) sub-populations (Anderson *et al*. 1991) is given by

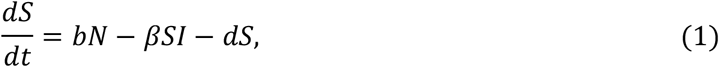

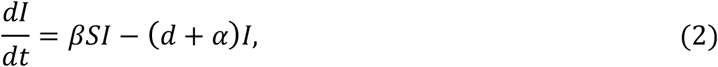

where *N*= *S*+*I, b* is the birth rate, *d* the baseline mortality rate, and *α* the parasite-induced mortality rate. The transmission coefficient *β*, while often written as a simple constant, actually incorporates both contact rate (*κ*) and infection probability given a contact (*ν*) (Begon *et al*. 2002), yielding

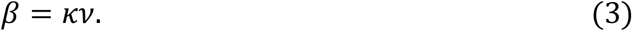

The infection probability *ν* can be further partitioned into the product of recipient susceptibility (*σ*) and donor infectiousness (*ι*),

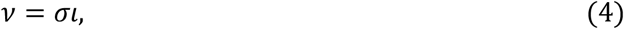

where *σ* and *ι* take values in [0,1], with higher values representing greater susceptibility or greater infectiousness, respectively. Thus, host heterogeneity in susceptibility and infectiousness can be incorporated by dividing the population into sub-populations comprising individuals that share the same values of susceptibility and infectiousness. Precisely how this is done depends on whether infectiousness is determined by a RD or DD scenario.

### Recipient dependence (RD)

Under this scenario recipient hosts are assigned to an infected sub-population based on a fixed, pre-determined trait inherent to that individual. Supposing that there are *n* unique pairs of trait values (*σ*_*j*_, *ι*_*j*_), 1 ≤ *j* ≤ *n*, we divide the population into *n* susceptible sub-populations *S*_*j*_, and *n* infected sub-populations *I*_*j*_, where the *j*th sub-populations share the *j*th trait pair. Thus, a RD heterogeneous analogue of Equations (1)-(2) is

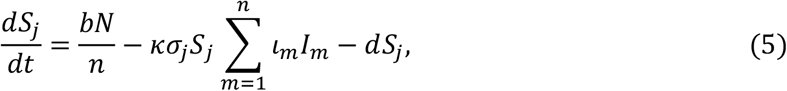

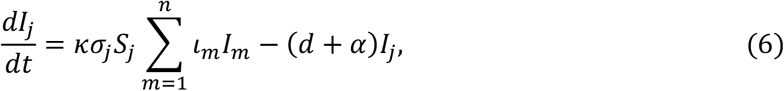

where we have additionally assumed that birth rate *b*, baseline mortality rate *d*, and parasite-induced mortality rate *α*, are equal across sub-populations. For the sake of simplicity, births are evenly distributed across susceptible sub-populations, effectively assuming non-inherited host heterogeneity, a phenomenon that has been previously observed, for example in *Daphnia magna* (Ben-Ami *et al*. 2008). We emphasise that in the RD scenario, the fixed traits of individuals determine the susceptible and infected sub-population to which they belong, so that all individuals in a susceptible sub-population move to the same infected sub-population upon becoming infected.

### Donor dependence (DD)

In this scenario, recipient hosts acquire their infectiousness trait when they become infected. Like in the RD scenario, we divide the susceptible sub-population into *n*_*S*_ sub-populations *S*_*j*_, each with susceptibility *σ*_*j*_, 1 ≤ *j* ≤ *n*_*S*_. Under the DD scenario, however, individuals are assigned the infectiousness of the donor host that infected them. We therefore define *I*_*j,k*_ to be those individuals with susceptibility *σ*_*j*_ that were infected by an individual with infectiousness *ι*_*k*_, 0 ≤ *k* ≤ *n*_*I*_, and so now share that same infectiousness value. Thus, a DD heterogeneous analogue of Equations (1)-(2) is

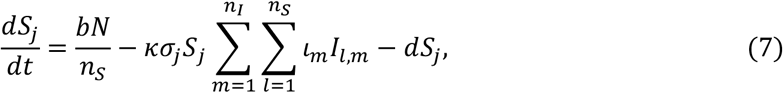

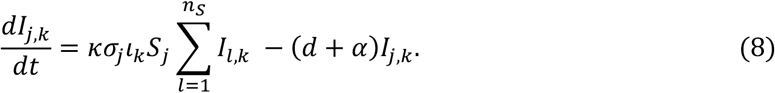

*b, d* and *α* are again assumed to be equal for all infected sub-populations. Unlike the RD model, the infected sub-population that the recipient host will join is not pre-determined before infection, so individuals in the same susceptible sub-population do not always move to the same infected sub-population upon infection. As such, the DD model has *n*_*S*_ susceptible sub-populations and *n*_*S*_*n*_*I*_ infectious sub-populations, whereas the RD model has equal numbers of both susceptible and infected sub-populations.

Equations (7)-(8), which will be useful when quantifying population-level heterogeneity, can be simplified by defining 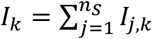, *i*.*e*., the sum of all individuals with the same infectiousness value, regardless of their initial susceptibility value. Summing Equation (8) over 0 ≤ *j* ≤ *n*_*S*_ yields

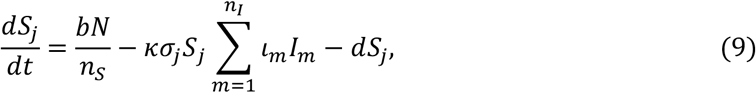

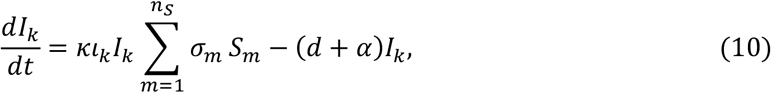

describing the dynamics of all infected individuals with infectiousness *ι*_(_. This simplified form of the system is useful for calculating *R*_*0*_. Note that we have used the same notation in the RD and the DD models to define similar but not precisely equivalent variables, and therefore we rely on context to provide clarity about which is being referred to throughout the remainder of this work.

### Quantifying population-level heterogeneity

We quantified population-level heterogeneity (hereafter referred to simply as ‘heterogeneity’) after (Laliberté *et al*. 2010), which is applicable to a wide range of systems and can deal with multiple traits and missing values (Olusoji *et al*. 2023). We undertook heterogeneity calculations in the context of two-dimensional *σ, ι* trait space, in which susceptibility (*σ*) and infectiousness (*ι*) form the two axes.

We first calculated the abundance-weighted centroid (*c*, hereafter referred to as the ‘centroid’) of the population, given simply as the population mean of each trait (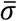 and 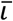, Figure 1A & C). We then calculated the heterogeneity score, *h*, by finding the mean abundance-weighted, Euclidean distance to the centroid (e.g. *z*_1_ in Figure 1B, *z*_11_ in Figure 1D) of all sub-populations. We calculated initial heterogeneity using the initial abundances of the sub-populations and final (equilibrium) heterogeneity using equilibrium sub-population abundances.

**Figure 1.**
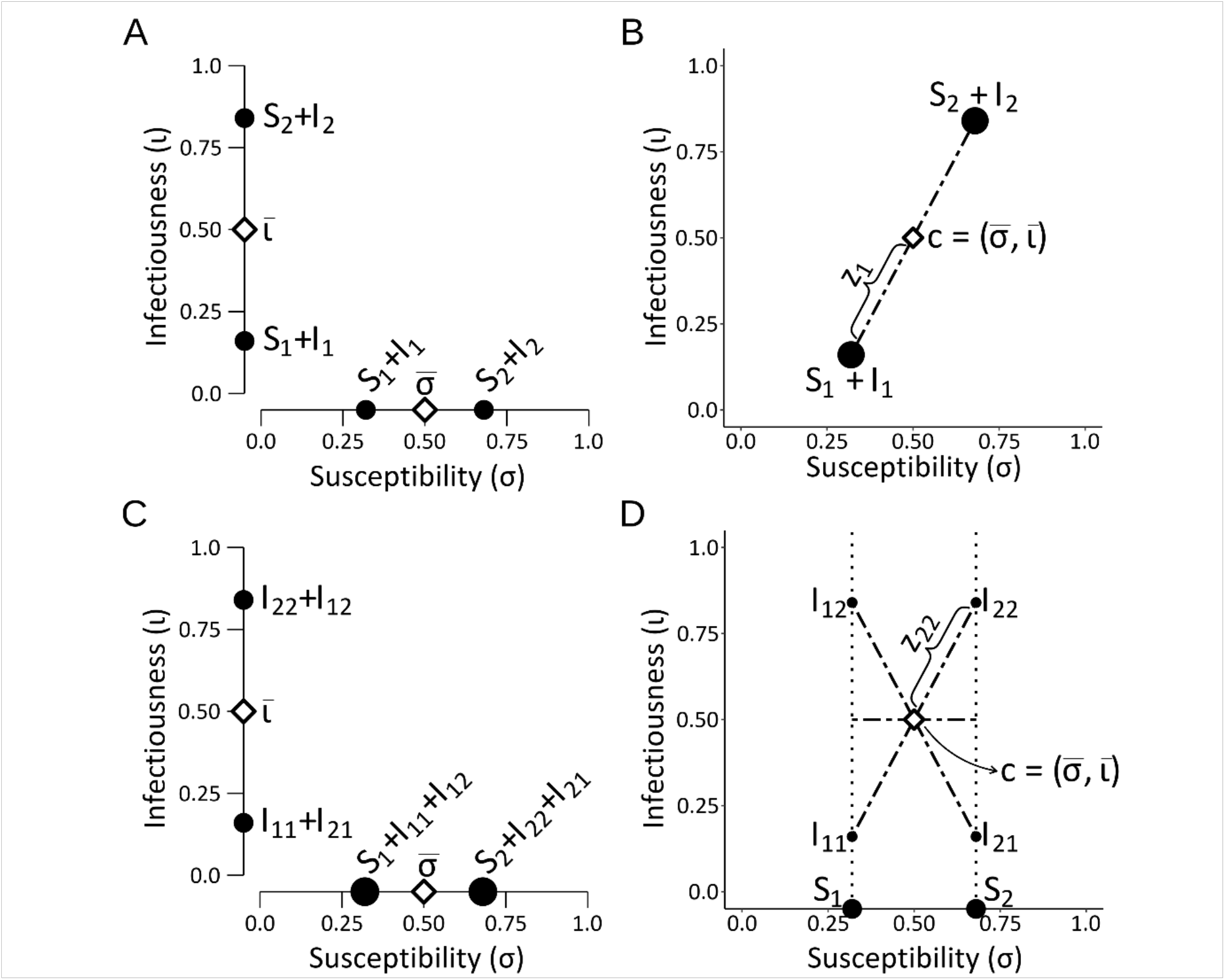
Schematic representation for calculating heterogeneity for the RD (A, B) and DD (C, D) scenarios. For both the RD and DD scenarios the population mean susceptibility 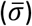 and population mean infectiousness 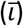 are calculated in a single dimension (A and C). The sizes of the black dots indicate the relative abundances of the relevant sub-populations. (B) Heterogeneity for the RD scenario is the mean Euclidean distance (*z*_*j*_), weighted by the abundance of each sub-population, to the centroid (*c*). (D) Heterogeneity for the DD scenario is the mean of the Euclidean distances (*z*_*j*(_) of the infected sub-populations to the centroid, and the single dimension distance (*σ*) of the susceptible sub-populations, weighted by the abundance of each sub-population; here the values for *S*_1_and *S*_2_ form lines rather than points in *σ, ι* trait space because they have no infectiousness values. In all panels the diameters of the black circles represent the abundance of the sub-population.

The calculation of heterogeneity differs between the RD and DD scenarios in how sub-populations were grouped, and abundances calculated. Specifically, for RD, all individuals in the same susceptible sub-population move to the same infected sub-population, and so we summed the abundances of the corresponding susceptible and infected sub-populations. These grouped sub-populations were then used to calculate *h*. For DD, each infected sub-population’s Euclidean distance to the centroid was calculated using both its *σ* and *ι* values (e.g. *I*_11_ in Figure 1D) while, because susceptible sub-populations only had a *σ* value, their distance to the centroid was calculated in a single dimension (e.g. *S*_1_ in Figure 1D), and so there were no grouped sub-populations used in calculating *h*. The formulae for heterogeneity calculations for both the RD and DD scenarios can be found in the Supplementary Information (SI).

### Model Analyses

We analysed both the RD and DD scenarios under three heterogeneity contexts:

#### 1) Bipartite heterogeneity

Where the population is divided into sub-populations corresponding to two distinct pairs of susceptibility and infectiousness values.

#### 2) *Tripartite isometric heterogeneity*

Where the population is divided into sub-populations corresponding to three distinct pairs of susceptibility and infectiousness values, equidistant from each other in *σ, ι* trait space.

#### 3) Tripartite non-isometric heterogeneity

Where the population is divided into sub-populations corresponding to three distinct pairs of susceptibility and infectiousness values, but which are not necessarily equidistant in *σ, ι* trait space.

Here we present analyses relating to context 1; contexts 2 and 3 are presented in the SI.

For the bipartite heterogeneity context, we set the initial number of susceptibles per sub-population to 49 and the number of infecteds per sub-population to 1 (RD, 2 sub-populations) or 0.5 (DD, 4 sub-populations). Initial mean population susceptibility and infectiousness trait values were 0.5; thus *σ*_2_= 1 − *σ*_1_ and *ι*_2_= 1 − *ι*_1_. All other parameter values were the same for all analyses (Table 1). By varying *σ*_1_ and *ι*_1_, we varied the initial population heterogeneity while maintaining the same initial mean population trait values. Hence, the effects of changing heterogeneity were decoupled from the effects of changing initial mean population trait values.

**Table 1.** Parameter definitions and values for all analyses of the RD and DD models.

| Model parameter | Definition | Value |
| --- | --- | --- |
| $\kappa$ | Contact rate per individual per time | 0.5 |
| $b$ | Birth rate per time | 1.5 |
| $d$ | Mortality rate per time | 1 |
| $\alpha$ | Parasite-induced mortality rate per time | 0.7 |

We generated 2,601 unique combinations of *σ* and *ι* trait values, and used these to analyse both the RD and DD models to understand how heterogeneity in susceptibility and infectiousness affected three key descriptors of epidemiological dynamics: (i) the basic reproduction number (*R*_*0*_), a measure of epidemic potential (Anderson *et al*. 1991); (ii) the equilibrium host population abundance, which quantifies the impact of the parasite on the host population; and (iii) the change in heterogeneity between the initial and final state of the system, indicating how heterogeneity changes with epidemic progression (presented in the SI).

### Calculating *R*_***0***_

*R*_*0*_ predicts the risk of an epidemic occurring, as well as the size and severity of that epidemic (Anderson *et al*. 1991; Heffernan *et al*. 2005), and also the effort needed to control and eliminate a parasite from a population (Roberts 2007). Thus, understanding the effect of heterogeneity on *R*_*0*_ provides considerable insight into how host heterogeneity in susceptibility and infectiousness affects parasite transmission through a population.

Using next generation matrices (Diekmann *et al*. 2010) (shown in full in the SI), we calculated *R*_*0*_ for the each scenario to be

**RD**:

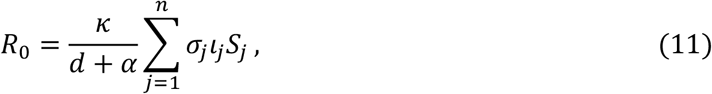

**DD:**

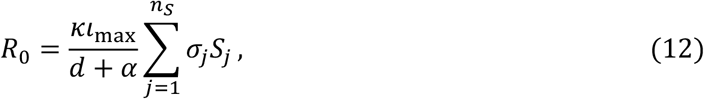

where *ι*_max_ is the maximum infectiousness value across all sub-populations.

### Equilibrium Analyses

When possible we calculated the equilibrium solutions of Equations (5)-(8) analytically. For parameter values for which this was not possible, we solved the system numerically over 50,000 time steps, which was sufficiently long to reach equilibrium. Any sub-population with susceptibility *σ*= 0 (*i*.*e*., completely resistant to infection) experiences unbounded growth, and so such cases were omitted from equilibrium analyses.

We also used these equilibrium solutions to calculate equilibrium population-level heterogeneity (following the method described above) to test whether the population-level heterogeneity changed with epidemiological progress.

### Software packages

Plots were generated in R (R Core Team 2022) using packages ggplot2 (Wickham 2016), ggforce (Pedersen 2022a), scales (Wickham *et al*. 2022), showtext (Qiu 2022), pBrackets (Schulz 2021), patchwork (Pedersen 2022b) and latex2exp (Meschiari 2022). Equilibrium analyses were conducted using Mathematica (Wolfram Research Inc. 2022). Heterogeneity and *R*_*0*_ values were calculated in R (R Core Team 2022).

## Results

Here we present analysis of the bipartite heterogeneity context, where there are two sub-populations. The results for the tripartite heterogeneities are broadly consistent with the bipartite analyses, highlighting the generality of the findings; we describe those results in the SI.

### (i) How heterogeneity affects *R*_*0*_ under RD v DD scenarios

#### Recipient dependence

In this scenario as heterogeneity increases *R*_*0*_ can increase (Figure 2A, red points), decrease (Figure 2A, blue points) or remain the same as the homogenous case (Figure 2A, yellow points). Whether it increases, decreases or remains the same depends on the population-level covariance between susceptibility and infectiousness. Mapping *R*_*0*_ values onto *σ, ι* trait space (Figure 2B) (where the sub-populations are mirrored across the centre point (*σ*_2_= 1 − *σ*_1_ and *ι*_2_= 1 − *ι*_1_)), shows that if there is heterogeneity in one trait only, *R*_*0*_ is the same as in the homogeneous population (*h*= 0) (Figure 2B yellow shading). This remains true even at high levels of heterogeneity. However, with positive covariance between susceptibility and infectiousness, *R*_*0*_ increases compared with the homogeneous population (Figure 2B, Supplementary Equation (S13)). *R*_*0*_ is at its maximum when one sub-population has maximal infectiousness and susceptibility values of 1, while the other sub-population has values of 0 for both; *i*.*e*. when the sub-populations lie at extremes of the positive diagonal in *σ, ι* trait space (Figure 2B). Conversely, with a negative covariance between susceptibility and infectiousness, *R*_*0*_ is lower than in the homogenous case (Figure 2B). In the extreme case, when one sub-population consists of only completely resistant hosts (susceptibility = 0, infectiousness = 1), and the other sub-population of completely dead-end hosts (susceptibility = 1, infectiousness = 0), such that they lie at extremes of the negative diagonal in *σ, ι* trait space, no individuals can both be infected and transmit onwards, resulting in an *R*_*0*_ of 0 (Figure 2B).

**Figure 2.**
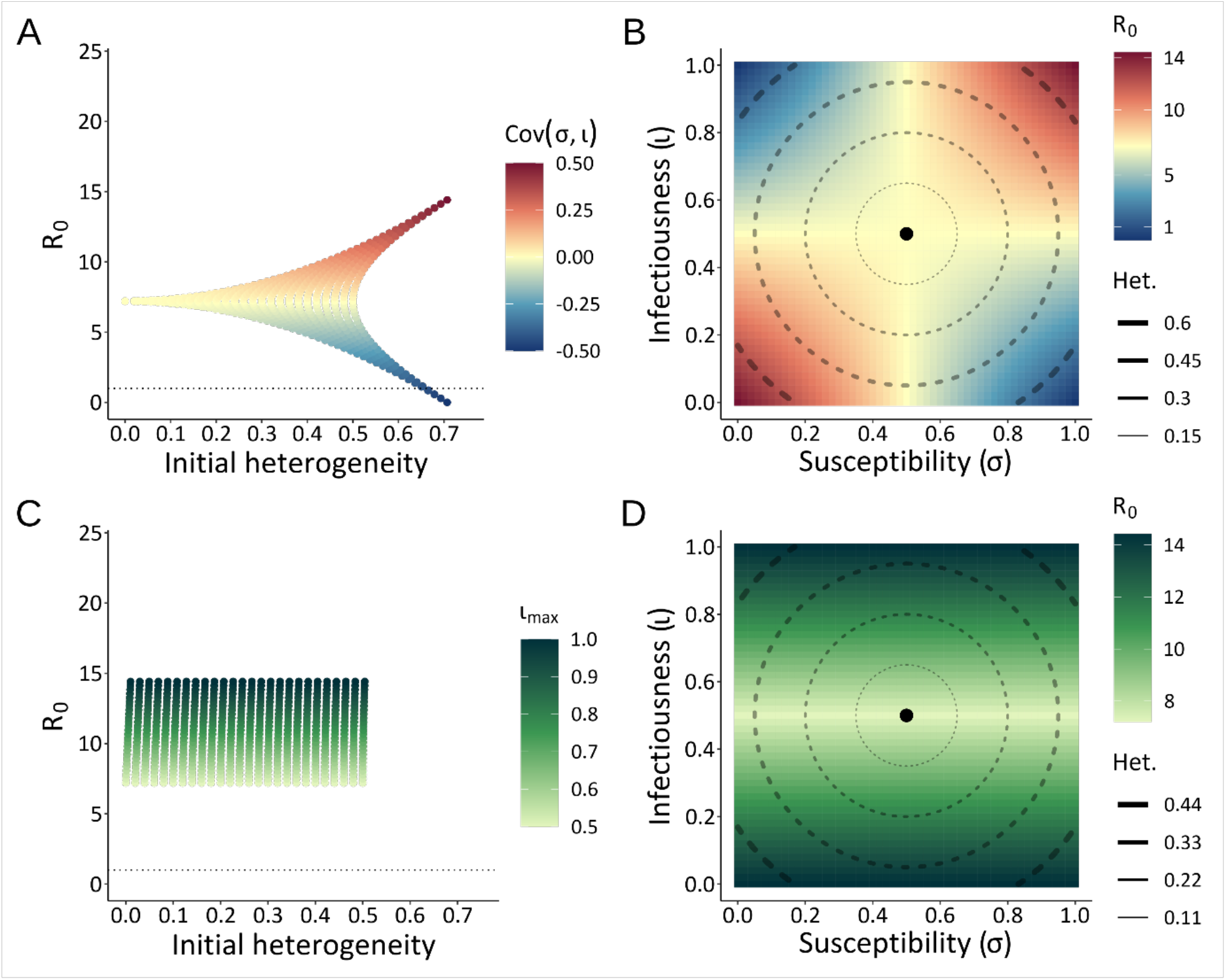
The effect of initial heterogeneity on *R*_*0*_ for the RD (A, B) and DD (C, D) scenarios. The dotted line in (A, C) is where *R*_*0*_ = 1. For RD (A) *R*_*0*_ changes as initial heterogeneity increases, and with the covariance between susceptibility (*σ*) and infectiousness (*ι*), as indicated by the colour bar. (B) shows *R*_*0*_ (the colour scale) plotted in *σ, ι* trait space with the centroid (*c*) at *σ*= *ι*= 0.5, and concentric dashed-line circles showing increasing heterogeneity; the positions of the sub-populations are mirrored across the centroid (*σ*_2_= 1 − *σ*_1_ and *ι*_2_= 1 − *ι*_1_). For DD (C) *R*_*0*_ does not change as initial heterogeneity increases, but scales with maximum infectiousness (*ι*_max_), as the green colour bar. (D) is the DD version of panel (B), but note that the *R*_*0*_ and heterogeneity scales differ between (B) and (D).

When there is a high degree of heterogeneity, there is a bifurcating distribution of points (Figure 2A) due to the boundaries of *σ, ι* trait space that restrict the number of viable trait combinations (*i*.*e*., where both susceptibility and infectiousness are between 0 and 1 for all sub-populations). For example, with maximum heterogeneity there are only two possible configurations of the sub-populations such that they lie at the opposite extremes of the diagonals in *σ, ι* trait space, resulting in just two points (Figure 2A).

In summary, increasing heterogeneity in the RD scenario can lead to increasingly divergent *R*_*0*_, compared to the homogeneous population. But, whether *R*_*0*_ is greater or less than the homogeneous population is determined by whether the population-level covariance between susceptibility and infectiousness is positive or negative, respectively. When the covariance is zero (heterogeneity in either susceptibility or infectiousness, but not both), *R*_*0*_ is the same as the homogeneous population, regardless of the degree of heterogeneity.

#### Donor dependence

In the DD scenario, *R*_*0*_ is affected in a qualitatively different fashion than in the RD scenario. Specifically, increasing heterogeneity does not lower *R*_*0*_ compared to the homogenous population (Figure 2C). Heterogeneity generally has little effect on *R*_*0*_, and changes in susceptibility alone do not affect *R*_*0*_, but changes in infectiousness do (Figure 2D).

In the DD scenario *R*_*0*_ is driven by the maximum infectiousness in the population (Figure 2C, Supplementary Equation (S13)). *R*_*0*_ is independent of susceptibility because, for equal sized susceptible sub-populations, there is a fixed mean value across the population (details in SI). Hence, *R*_*0*_ changes only along the infectiousness axis (Figure 2D), not along the susceptibility axis. For the same overall degree of heterogeneity (*i*.*e*., vertical slice in Figure 2C), increasing heterogeneity in infectiousness (and a necessary decrease in heterogeneity in susceptibility) increases *R*_*0*_. Susceptibility can only affect *R*_*0*_ when the initial susceptible sub-population abundances are not equal (see SI).

#### Summary of effects of heterogeneity on R_**0**_

In both the RD and DD scenarios increasing heterogeneity can lead to increasingly divergent *R*_*0*_ values compared to the homogeneous case. But, what drives this effect on *R*_*0*_ differs between the two scenarios. In the RD scenario it is the population-level covariance between susceptibility and infectiousness; in the DD scenario it is the maximum infectiousness in the population.

### (ii) How heterogeneity affects host abundance under RD v DD scenarios

Equilibrium total host abundance generally increases with increasing initial heterogeneity, as does the variability in equilibrium abundance, for both the RD and DD scenarios (Figure 3). However, specific aspects of these relationships differ between the two scenarios.

**Figure 3.**
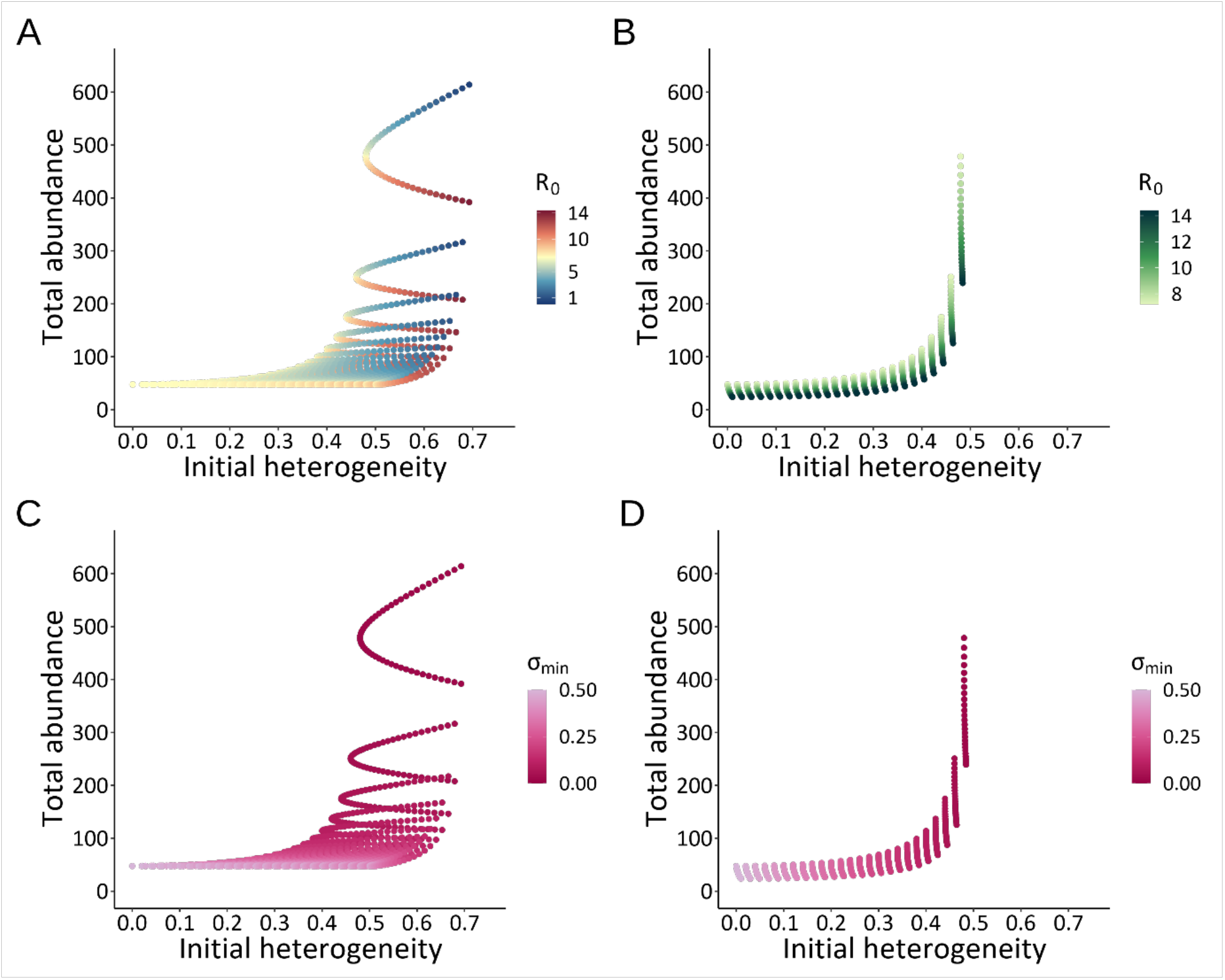
The effect of initial heterogeneity on equilibrium total host abundance for RD (A, C) and DD (B, D) scenarios. (A) and (C) are the same RD plots, but differently coloured, (A) for *R*_*0*_ and (C) for minimum population-level susceptibility (*σ*_*min*_). (B) and (D) are the same DD plots, but differently coloured, (B) for *R*_*0*_ and (D) for *σ*_*min*_. Note that the *R*_*0*_ scales differ between (A) and (B). An enlarged version of (A) and (C) can be found in the SI (Figure S2).

#### Recipient dependence

In the RD scenario there is a complex relationship between initial heterogeneity and host equilibrium abundance (Figure 3A & C, and see Figure S2 for an enlarged version at the lower host abundance range). Equilibrium abundances are grouped into parabolic ‘clusters’ where each cluster has equilibrium abundances that both increase and decrease from an initial baseline abundance as heterogeneity increases. Each cluster is determined by the minimum susceptibility in the population; all points in a cluster have the same population-level minimum susceptibility value (and so also the same maximum susceptibility value) (Figure 3C). Clusters are ordered by these minimum susceptibility values; those with the lowest minimum susceptibility values (*σ*_*min*_, Figure 3C) have the highest abundances. This is because in populations with low minimum susceptibility fewer individuals become infected, so that fewer hosts are exposed to parasite-induced mortality (*α*), which thereby increases overall host abundance.

The within-cluster divergence in host abundance that occurs with increasing heterogeneity arises due to increasingly divergent infectiousness values in the population. Since all points within a cluster have the same pair of *σ* values for the component sub-populations, an increase in heterogeneity is achieved by divergence in the two infectiousness values. This divergence, and so increase in heterogeneity, leads to changes in *R*_*0*_ within a cluster that changes equilibrium host abundance. High *R*_*0*_ values lead to lower abundances (the lower red tail of a cluster in Figure 3A), while low *R*_*0*_ values lead to higher abundances (the upper blue tail of a cluster in Figure 3A).

#### Donor dependence

In the DD scenario there are also clusters in equilibrium host abundance determined by the minimum susceptibility value (*σ*_*min*_) within the population (Figure 3D), but the points within each cluster are arranged in near-vertical lines, suggesting that heterogeneity has little effect on equilibrium host abundance within a cluster. As with the RD scenario, equilibrium host abundance is maximised when *R*_*0*_ is minimised (Figure 3B) which, for the DD scenario occurs when the maximum infectiousness in the population (*ι*_*max*_) is at its lowest (Figure 2C). Hence within each cluster, increasing maximum infectiousness increases *R*_*0*_, reducing equilibrium host abundance. The DD scenario generally shows lower equilibrium host abundances than the RD scenario because the minimum *R*_*0*_ values for the RD scenario are lower than the DD scenario.

## Discussion

While existing theory shows that host heterogeneity in susceptibility and infectiousness can affect population-level parasite transmission (*e*.*g*. Lloyd *et al*. (2020)), our findings demonstrate that these effects differ considerably depending how host infectiousness is determined. We show that the contrasting RD and DD scenarios result in different relationships between host heterogeneity in susceptibility and infectiousness and epidemiological outcomes, specifically *R*_*0*_ and host abundance.

While we find that *R*_*0*_ changes with increasing heterogeneity in both scenarios, there is a notable contrast between the two scenarios in what drives these effects, and the direction of the effects. The RD scenario shows divergent *R*_*0*_ values as heterogeneity increases, which is determined by the covariance between susceptibility and infectiousness. This finding is consistent with previous modelling: three models of vector-borne infections, where infectiousness was an inherent host trait (*i*.*e*. implicitly assuming RD), incorporated host heterogeneity in susceptibility and infectiousness and found that positive covariance between heterogeneities led to an increase in *R*_*0*_ relative to the homogeneous case, while negative covariance led to a decrease (Dietz 1980; Koella 1991; Vazquez-Prokopec *et al*. 2016). Thus, these models showed that in a RD scenario, population-level covariance between susceptibility and infectiousness determines *R*_*0*_, consistent with our findings.

In contrast, we find that in the DD scenario *R*_*0*_ is determined by the maximum infectiousness in the population, such that *R*_*0*_ increases as heterogeneity in infectiousness increases. Heterogeneity in susceptibility is largely irrelevant for the DD scenario, only becoming so if abundances are very different between sub-populations. There are fewer other studies that consider DD-like scenarios, but one example is the hypothesis that patterns of SARS-CoV-2 transmission may be due to superspreaders tending to generate new superspreaders, for example through a dose-dependent effect (Beldomenico 2020). A model exploring how *R*_*0*_ responded to this scenario (compared to the null model where superspreaders appeared randomly) showed that *R*_*0*_ increases with an increase in the probability that a superspreader generates additional superspreaders (Wanelik *et al*. 2023). Thus, moving from the null model to a DD-like scenario increased *R*_*0*_, suggesting that the DD scenario tends to increase *R*_*0*_, consistent with our findings.

The RD and DD scenarios also differ in how host heterogeneity affects equilibrium total host abundance. While host abundance tends to increase with increasing heterogeneity in both scenarios, the effect is more pronounced in the RD scenario than in the DD scenario. Further, for a given susceptibility value, RD and DD differ by generating divergent (RD) or monotonic (DD) changes in equilibrium host abundance, a pattern that becomes more pronounced with increasing heterogeneity.

These RD and DD scenarios are phenomenological characterisations that each encompass a variety of real-world processes. Empirical examples matching assumptions of the RD scenario include the finding that canaries’ nutritional status can affect their infectiousness with avian malaria (Cornet *et al*. 2014); rabbit myxoma virus infection status determines its infectiousness for co-infecting nematodes (Cattadori *et al*. 2007); several examples of different host strains exhibiting varying levels of infectiousness when infected with the same parasite isolates (Bolas-Fernandez *et al*. 1989; Jørgensen *et al*. 1998; Dorfman *et al*. 2024); a genetic basis for observed variation in host infectiousness in *Scophthalmus maximus* (Turbot) infected with a ciliate parasite (Anacleto *et al*. 2019). In addition, critical to understanding parasite transmission in a RD scenario, is covariance between susceptibility and infectiousness, and there are empirical examples of this. For example, it has been shown that juvenile *Cygnus columbianus bewickii* (Bewick’s swan) are more susceptible to avian influenza virus than adults and also have higher shedding rates than adults, demonstrating positive covariance between host susceptibility and infectiousness (Hoye *et al*. 2012). Alternatively, negative covariance between host susceptibility and infectiousness has been proposed as a mechanism for social insects avoiding parasitism by investing more in anti-parasitic behaviours to reduce their infectiousness in a colony with high susceptibility (Cremer *et al*. 2007), such that a highly susceptible colony would be less infectious than a less susceptible colony. At the individual level, negative covariance might occur when highly susceptible individuals acquire high levels of disease, thus increasing sickness behaviours that reduce host infectiousness (Araujo *et al*. 2016).

Empirical examples matching assumptions of the DD scenario are calves infected with three different doses of bovine viral diarrhoea virus (BVDV), where the most infectious individuals were those given the highest viral dose, due to a longer infectious period (Strong *et al*. 2015). Similar patterns have been found with a number of other host-parasite systems (Gaskell *et al*. 1979; Mumford *et al*. 1990; Zarkov 2012).

In reality, many host-parasite systems are unlikely to fit solely under either RD or DD scenarios, instead falling somewhere between the two. For example, in the BVDV calf infection study, while calves infected with the highest BVDV doses were the most infectious, there was still within-dose group heterogeneity in infectiousness (Strong *et al*. 2015), which may have been caused by traits inherent to individual calves. Thus, while the overall system may be best described by the DD scenario, there are still aspects of the RD scenario at play. The reverse can also be true. For example, although myxoma-infected rabbits may be more nematode infectious (Cattadori *et al*. 2007), aligning with the RD scenario, there may still be some DD involved. Specifically, the nematode transmits between hosts via a faecal-oral pathway (Cattadori *et al*. 2007). Therefore, a rabbit might become more infectious when infected by a host with a high parasite burden, because it will acquire more parasites in turn. These examples demonstrate that *R*_*0*_ will often be affected by both the covariance between susceptibility and infectiousness (RD) as well as the maximum infectiousness in the population (DD), though which of these two measures is more influential will depend on where on the spectrum of RD to DD that specific host-parasite system exists.

Previous work has typically treated how a host’s infectiousness is determined as a black box, generally assuming it is a fixed property of the recipient host. This overlooks the importance of how a host’s infectiousness is determined in influencing the consequences of host heterogeneity for parasite transmission. Yet this process can have real-world consequences. For instance, there is interest in breeding parasite resistant livestock to reduce the substantial economic and climatic costs caused by parasites in livestock systems (Knap *et al*. 2020). However, to determine the livestock traits that are most appropriate to select, it will be important to consider where the host-parasite system falls on the RD-DD spectrum. For example, in a RD scenario, selecting for reduced parasite susceptibility (*i*.*e*., for resistance), versus in a DD scenario selecting for reduced parasite shedding. The work we present here shows the importance of explicitly considering how infectiousness is determined, and that ignoring it can lead to an incomplete understanding of the effects of host heterogeneities on parasite transmission. A failure to do so could have consequences for both future theoretical and empirical work.

## Acknowledgements

JAC was funded by a PhD studentship from the ‘Adapting to the Challenges of a Changing Environment (ACCE)’ Doctoral Training Partnership funded by the Natural Environment Research Council (NERC) (Grant NE/S00713X/1). ADD was funded by a NERC Pushing the Frontiers grant (Grant NE/X01424X/1).

## References

Adelman, J.S., Moyers, S.C., Farine, D.R. & Hawley, D.M. (2015) Feeder use predicts both acquisition and transmission of a contagious pathogen in a North American songbird. Proceedings of the Royal Society B: Biological Sciences, 282, 20151429.

Anacleto, O., Cabaleiro, S., Villanueva, B., Saura, M., Houston, R.D., Woolliams, J.A. & Doeschl-Wilson, A.B. (2019) Genetic differences in host infectivity affect disease spread and survival in epidemics. Scientific Reports, 9, 4924.

Anderson, R.M. & May, R.M. (1991) Infectious Diseases of Humans. Oxford University Press.

Araujo, A., Kirschman, L. & Warne, R.W. (2016) Behavioural phenotypes predict disease susceptibility and infectiousness. Biology Letters, 12, 20160480.

Barlow, N.D. (2000) Non-linear transmission and simple models for bovine tuberculosis. journal of animal ecology, 69, 703–713.

Begon, M., Bennett, M., Bowers, R.G., French, N.P., Hazel, S. & Turner, J. (2002) A clarification of transmission terms in host-microparasite models: numbers, densities and areas. Epidemiology & Infection, 129, 147–153.

Beldomenico, P.M. (2020) Do superspreaders generate new superspreaders? A hypothesis to explain the propagation pattern of COVID-19. International Journal of Infectious Diseases, 96, 461–463.

Ben-Ami, F., Regoes, R.R. & Ebert, D. (2008) A quantitative test of the relationship between parasite dose and infection probability across different host–parasite combinations. Proceedings of the Royal Society B: Biological Sciences, 275, 853–859.

Bolas-Fernandez, F. & Wakelin, D. (1989) Infectivity of Trichinella isolates in mice is determined by host immune responsiveness. Parasitology, 99, 83–88.

Brault, A.C., Langevin, S.A., Bowen, R.A., Panella, N.A., Biggerstaff, B.J., Miller, B.R. & Komar, N. (2004) Differential virulence of West Nile strains for American crows. Emerging infectious diseases, 10, 2161.

Cattadori, I.M., Albert, R. & Boag, B. (2007) Variation in host susceptibility and infectiousness generated by co-infection: the myxoma–Trichostrongylus retortaeformis case in wild rabbits. Journal of The Royal Society Interface, 4, 831–840.

Cornet, S., Bichet, C., Larcombe, S., Faivre, B. & Sorci, G. (2014) Impact of host nutritional status on infection dynamics and parasite virulence in a bird-malaria system. journal of animal ecology, 83, 256–265.

Coutinho, F.A.B., Massad, E., Lopez, L.F., Burattini, M.N., Struchiner, C.J. & Azevedo-Neto, R.S.d. (1999) Modelling heterogeneities in individual frailties in epidemic models. Mathematical and computer modelling, 30, 97–115.

Cremer, S., Armitage, S.A. & Schmid-Hempel, P. (2007) Social immunity. Current biology, 17, R693–R702.

Dai, Y., Liu, M., Cheng, X., Shen, X., Wei, Y., Zhou, S., Yu, S. & Ding, C. (2013) Infectivity and pathogenicity of Newcastle disease virus strains of different avian origin and different virulence for mallard ducklings. Avian Diseases, 57, 8–14.

Diekmann, O., Heesterbeek, J. & Roberts, M.G. (2010) The construction of next-generation matrices for compartmental epidemic models. Journal of The Royal Society Interface, 7, 873–885.

Dietz, K. (1980) Models for vector-borne parasitic diseases. Vito Volterra Symposium on Mathematical Models in Biology, pp. 264–277. Springer.

Dorfman, B., Marcos-Hadad, E., Tadmor-Levi, R. & David, L. (2024) Disease resistance and infectivity of virus susceptible and resistant common carp strains. Scientific Reports, 14, 4677.

Dwyer, G., Elkinton, J.S. & Buonaccorsi, J.P. (1997) Host heterogeneity in susceptibility and disease dynamics: tests of a mathematical model. The American Naturalist, 150, 685–707.

Gaskell, R. & Povey, R. (1979) The dose response of cats to experimental infection with feline viral rhinotracheitis virus. Journal of Comparative Pathology, 89, 179–191.

Heffernan, J.M., Smith, R.J. & Wahl, L.M. (2005) Perspectives on the basic reproductive ratio. Journal of The Royal Society Interface, 2, 281–293.

Hickson, R. & Roberts, M. (2014) How population heterogeneity in susceptibility and infectivity influences epidemic dynamics. Journal of Theoretical Biology, 350, 70–80.

Hoye, B.J., Fouchier, R.A. & Klaassen, M. (2012) Host behaviour and physiology underpin individual variation in avian influenza virus infection in migratory Bewick’s swans. Proceedings of the Royal Society B: Biological Sciences, 279, 529–534.

Jørgensen, L., Leathwick, D., Charleston, W., Godfrey, P., Vlassoff, A. & Sutherland, I. (1998) Variation between hosts in the developmental success of the free-living stages of trichostrongyle infections of sheep. International Journal for Parasitology, 28, 1347–1352.

Knap, P.W. & Doeschl-Wilson, A. (2020) Why breed disease-resilient livestock, and how? Genetics Selection Evolution, 52, 1–18.

Koella, J.C. (1991) On the use of mathematical models of malaria transmission. Acta tropica, 49, 1–25.

Kong, L., Wang, J., Han, W. & Cao, Z. (2016) Modeling heterogeneity in direct infectious disease transmission in a compartmental model. International journal of environmental research and public health, 13, 253.

Laliberté, E. & Legendre, P. (2010) A distance-based framework for measuring functional diversity from multiple traits. Ecology, 91, 299–305.

Lemieux, J.E., Siddle, K.J., Shaw, B.M., Loreth, C., Schaffner, S.F., Gladden-Young, A., Adams, G., Fink, T., Tomkins-Tinch, C.H. & Krasilnikova, L.A. (2021) Phylogenetic analysis of SARS-CoV-2 in Boston highlights the impact of superspreading events. Science, 371, eabe3261.

Lloyd, A.L., Kitron, U., Perkins, T.A., Vazquez-Prokopec, G.M. & Waller, L.A. (2020) The basic reproductive number for disease systems with multiple coupled heterogeneities. Mathematical biosciences, 321, 108294.

Lloyd-Smith, J.O., Schreiber, S.J., Kopp, P.E. & Getz, W.M. (2005) Superspreading and the effect of individual variation on disease emergence. Nature, 438, 355–359.

Mantzaris, A. & Higham, D. (2013) Dynamic communicatbility predicts infectiousness. Temporal Networks (eds P. Holme & J. Saramäki). Springer, Berlin, Heidelberg.

Matthews, L., McKendrick, I.J., Ternent, H., Gunn, G., Synge, B. & Woolhouse, M. (2006) Super-shedding cattle and the transmission dynamics of Escherichia coli O157. Epidemiology & Infection, 134, 131–142.

McCallum, H., Fenton, A., Hudson, P.J., Lee, B., Levick, B., Norman, R., Perkins, S.E., Viney, M., Wilson, A.J. & Lello, J. (2017) Breaking beta: deconstructing the parasite transmission function. Philosophical Transactions of the Royal Society B: Biological Sciences, 372, 20160084.

Medina, L., Castillo, C., Liempi, A., Herbach, M., Cabrera, G., Valenzuela, L., Galanti, N., de los Angeles Curto, M., Schijman, A.G. & Kemmerling, U. (2018) Differential infectivity of two Trypanosoma cruzi strains in placental cells and tissue. Acta tropica, 186, 35–40.

Meschiari, S. (2022) latex2exp: Use LaTex Expression in Plots.

Miller, J.C. (2007) Epidemic size and probability in populations with heterogeneous infectivity and susceptibility. Physical Review E, 76, 010101.

Mumford, J.A., Hannant, D. & Jessett, D. (1990) Experimental infection of ponies with equine influenza (H3N8) viruses by intranasal inoculation or exposure to aerosols. Equine Veterinary Journal, 22, 93–98.

Olusoji, O.D., Barabás, G., Spaak, J.W., Fontana, S., Neyens, T., De Laender, F. & Aerts, M. (2023) Measuring individual-level trait diversity: a critical assessment of methods. Oikos, 2023, e09178.

Pedersen, T.L. (2022a) ggforce: Accelerating ‘ggplot2’.

Pedersen, T.L. (2022b) patchwork: The Composer of Plots.

Qiu, Y. (2022) showtext: Using Fonts More Easily in R Graphs.

R Core Team (2022) R: A language and environment for statistical computing. R Foundation for Statistical Computing, Vienna, Austria.

Roberts, M. (2007) The pluses and minuses of R0. Journal of The Royal Society Interface, 4, 949–961.

Schulz, A. (2021) pBrackets: Plot Brackets.

Stephenson, J.F., Young, K.A., Fox, J., Jokela, J., Cable, J. & Perkins, S.E. (2017) Host heterogeneity affects both parasite transmission to and fitness on subsequent hosts. Philosophical Transactions of the Royal Society B: Biological Sciences, 372, 20160093.

Streicker, D.G., Fenton, A. & Pedersen, A.B. (2013) Differential sources of host species heterogeneity influence the transmission and control of multihost parasites. Ecology letters, 16, 975–984.

Strong, R., La Rocca, S.A., Paton, D., Bensaude, E., Sandvik, T., Davis, L., Turner, J., Drew, T., Raue, R. & Vangeel, I. (2015) Viral dose and immunosuppression modulate the progression of acute BVDV-1 infection in calves: evidence of long term persistence after intra-nasal infection. PLoS One, 10, e0124689.

VanderWaal, K.L. & Ezenwa, V.O. (2016) Heterogeneity in pathogen transmission: mechanisms and methodology. Functional Ecology, 30, 1606–1622.

Vazquez-Prokopec, G.M., Perkins, T.A., Waller, L.A., Lloyd, A.L., Reiner Jr, R.C., Scott, T.W. & Kitron, U. (2016) Coupled heterogeneities and their impact on parasite transmission and control. Trends in parasitology, 32, 356–367.

Wanelik, K.M., Begon, M., Fenton, A., Norman, R.A. & Beldomenico, P.M. (2023) Positive feedback loops exacerbate the influence of superspreaders in disease transmission. iScience, 26.

White, L.A., Forester, J.D. & Craft, M.E. (2018) Covariation between the physiological and behavioral components of pathogen transmission: host heterogeneity determines epidemic outcomes. Oikos, 127, 538–552.

Wickham, H. (2016) ggplot2: Elegant Graphics for Data Analysis. Springer-Verlag New York.

Wickham, H. & Seidel, D. (2022) scales: Scale Functions for Visualization.

Wolfram Research Inc. (2022) Mathematica. Wolfram Research Inc., Champaign, Illinois.

Woolhouse, M.E., Dye, C., Etard, J.-F., Smith, T., Charlwood, J., Garnett, G., Hagan, P., Hii, J., Ndhlovu, P. & Quinnell, R. (1997) Heterogeneities in the transmission of infectious agents: implications for the design of control programs. Proceedings of the National Academy of Sciences, 94, 338–342.

Yates, A., Antia, R. & Regoes, R.R. (2006) How do pathogen evolution and host heterogeneity interact in disease emergence? Proceedings of the Royal Society B: Biological Sciences, 273, 3075–3083.

Zarkov, I. (2012) The importance of inoculation dose of avian H6N2 influenza a virus on virus shedding of Anas plathyrynchos ducks after induced generalised infection. Trakia Journal of Sciences, 10, 58–61.

