## Supplementary Information for "The consequences of host heterogeneity for parasite transmission depend on how host infectiousness is determined"

### 1. Calculation of heterogeneity scores

We adapted the method outlined in (Laliberté & Legendre 2010) to calculate heterogeneity as the weighted mean of the Euclidean distances in trait space of each sub-population from the population mean, where the weights are given by the abundances of each sub-population. The following calculations are presented graphically in Figure 1 for the bipartite heterogeneity context $n = 2$, $\sigma_2 = 1 - \sigma_1$ and $\iota_2 = 1 - \iota_1$.

***Recipient dependence***

In the RD scenario, there are $n$ susceptible and $n$ infected sub-populations, with the $j$th sub-populations sharing the same values of $\sigma$ and $\iota$. The population mean trait values are therefore

$$\bar{\sigma} = \frac{1}{N}\sum_{j=1}^{n} \sigma_j (S_j + I_j), \quad (S1)$$

$$\bar{\iota} = \frac{1}{N}\sum_{j=1}^{n} \iota_j \left(S_j + I_j\right). \quad (S2)$$

The Euclidean distance of the $j$th sub-population to the population mean is

$$z_j = \sqrt{\left(\bar{\sigma} - \sigma_j\right)^2 + \left(\bar{\iota} - \iota_j\right)^2}, \quad (S3)$$

and so population heterogeneity is given by

$$h = \frac{1}{N}\sum_{j=1}^{n} z_j (S_j + I_j)\,. \quad (S4)$$

***Donor dependence***

In the DD scenario, there are $n_S$ susceptible sub-populations, each with no infectiousness trait, and $n_S n_I$ infected sub-populations, each with both traits. The population mean trait values are therefore

$$\bar{\sigma} = \frac{1}{N}\sum_{j=1}^{n_S} \sigma_j \left( S_j + \sum_{k=1}^{n_I} I_{j,k} \right), \qquad (S5)$$

$$\bar{\iota} = \frac{1}{\sum_{k=1}^{n_I}\sum_{j=1}^{n_S} I_{j,k}} \sum_{k=1}^{n_I}\sum_{j=1}^{n_S} \iota_k \, I_{j,k} \,. \qquad (S6)$$

Note that the population size for calculating the mean infectiousness is the sum of all infected sub-populations, not $N$. The Euclidean distance for the susceptible sub-populations in this case is simply $z_j = \left|\bar{\sigma} - \sigma_j\right|$ (see Figure 1D), while for the infectious sub-populations it is

$$z_{j,k} = \sqrt{\left(\bar{\sigma} - \sigma_j\right)^2 + (\bar{\iota} - \iota_k)^2}. \qquad (S7)$$

Hence population heterogeneity is given by

$$h = \frac{1}{N}\left( \sum_{j=1}^{n_S} \left( z_j S_j + \sum_{k}^{n_I} z_{j,k} I_{j,k} \right) \right). \qquad (S8)$$

### 2. $R_0$ calculation through next generation matrices

To calculate $R_0$, we followed the method described in (Diekmann *et al.* 2010), in which $R_0$ is given by the dominant eigenvalue of the next-generation matrix $K = -T\Sigma^{-1}$, where $T$ comprises the rates of the infection transmission events in the infected subsystem, and $\Sigma$ the rates of the transition (e.g. mortality) events.

***Recipient dependence***

In the RD scenario, the infected subsystem is Equation (6). In this subsystem, the element in the $x$th row and $y$th column of the transmission matrix is

$$T_{xy} = \kappa \sigma_x S_x \iota_y, \quad 1 \le x \le n, 1 \le y \le n, \qquad (S9)$$

and the transition matrix is $\Sigma = -(d+\alpha)I_n$, where $I_n$ is the $n$-dimensional identity matrix. Thus, the elements of the next-generation matrix are

$$K_{xy} = \frac{\kappa \sigma_x S_x \iota_y}{d+\alpha}, \quad 1 \le x \le n, 1 \le y \le n. \qquad (S10)$$

Now, $K$ has rank 1, because each row is a scalar multiple of $(\iota_1, \dots, \iota_n)$, and therefore has precisely one non-zero eigenvalue. Because the sum of the eigenvalues of a matrix is equal to its trace, we therefore have

$$R_0 = \frac{\kappa}{d+\alpha} \sum_{j=1}^{n} \sigma_j \iota_j S_j \,. \qquad (S11)$$

The covariance of the two trait values is

$$\operatorname{cov}(\sigma, \iota) = \frac{1}{N} \sum_{j}^{n} \sigma_j \iota_j \left(S_j + I_j\right) - \bar{\sigma}\,\bar{\iota}; \qquad (S12)$$

hence, if $S_j \gg I_j$, as at the beginning of an infection, we can write

$$R_0 \approx \frac{\kappa N}{d+\alpha} (\operatorname{cov}(\sigma, \iota) + \bar{\sigma}\,\bar{\iota}). \qquad (S13)$$

In other words, $R_0$ scales with the covariance of $\sigma$ and $\iota$, as seen in Figure 2 for $n = 2$.

***Donor dependence***

In the DD scenario, the infected subsystem is Equation (10); note we use the system as written in terms of $I_k$, not $I_{j,k}$ as the susceptibility of infected classes does not impact parasite transmission. In this subsystem, the transmission matrix is given by

$$T = \kappa \sum_{j=1}^{n_S} \sigma_j S_j \operatorname{diag}(\boldsymbol{\iota}_{DD}), \qquad (S14)$$

where $\boldsymbol{\iota}_{DD} = (\iota_1, \dots, \iota_{n_I})$ is the vector of infectiousness values and $\mathrm{diag}(\boldsymbol{x})$ is the diagonal matrix with $\boldsymbol{x}$ on the diagonal and zeroes elsewhere. The transition matrix is $\Sigma = -(d+\alpha)I_{n_I}$, and so the next-generation matrix is

$$K = \frac{\kappa}{d+\alpha}\sum_{j=1}^{n_S} \sigma_j S_j \ \mathrm{diag}(\boldsymbol{\iota}_{DD}). \qquad (S15)$$

The eigenvalues of a diagonal matrix are simply its diagonal entries, and so in this case the maximal eigenvalue is

$$R_0 = \frac{\kappa \iota_{\max}}{d+\alpha}\sum_{j=1}^{n_S} \sigma_j S_j\,, \qquad (S16)$$

where $\iota_{\max}$ is maximum infectiousness value present in the system. In Figure 2 (where $n = 2$) we have equal abundances for susceptible sub-populations and $\sigma_2 = 1 - \sigma_1$, so it is $\iota_{\max}$ alone that determines how $R_0$ varies in trait space.

**3. $R_0$ with bipartite heterogeneity**

In this section, we present a mathematical analysis of $R_0$ when there two different values of susceptibility and infectiousness, *i.e.* $n = 2$ in RD and $n_S = n_I = 2$ in DD.

***Recipient dependence***

In this case, we define $N_j = S_j + I_j, j = 1{,}2$. From Equation (S4), we can therefore write population heterogeneity as

$$h = \frac{N_1}{N}\sqrt{(\bar{\sigma} - \sigma_1)^2 + (\bar{\iota} - \iota_1)^2} + \frac{N_2}{N}\sqrt{(\bar{\sigma} - \sigma_2)^2 + (\bar{\iota} - \iota_2)^2}. \qquad (S17)$$

Using the definition of trait means (Equations (S1)-(S2)), this can be rewritten as

$$h = \frac{N_1}{N}\sqrt{(\bar{\sigma} - \sigma_1)^2 + (\bar{\iota} - \iota_1)^2} + \frac{N_2}{N}\sqrt{\left(\bar{\sigma} - \frac{\bar{\sigma}N - \sigma_1 N_1}{N_2}\right)^2 + \left(\bar{\iota} - \frac{\bar{\iota}N - \iota_1 N_1}{N_2}\right)^2}$$

$$= \frac{N_1}{N}\sqrt{(\bar{\sigma} - \sigma_1)^2 + (\bar{\iota} - \iota_1)^2} + \frac{1}{N}\sqrt{(-\bar{\sigma}N_1 + \sigma_1 N_1)^2 + (-\bar{\iota}N_1 + \iota_1 N_1)^2}$$

$$= \frac{2N_1}{N}\sqrt{(\bar{\sigma} - \sigma_1)^2 + (\bar{\iota} - \iota_1)^2}. \qquad (S18)$$

Thus $N_1 z_1 = N_2 z_2$, and Equation (S18) describes a circle in $\sigma_1, \iota_1$ space, centred at trait means and with radius $hN/2N_1$. We therefore define the polar coordinates

$$\sigma_1 = \bar{\sigma} + \frac{\rho N}{2N_1}\cos\theta\,, \qquad (S19)$$

$$\iota_1 = \bar{\iota} + \frac{\rho N}{2N_1}\sin\theta, \qquad (S20)$$

and fix trait means at $\bar{\sigma}$ and $\bar{\iota}$. Hence, Equations (S1)-(S2) yield

$$\sigma_2 = \bar{\sigma} - \frac{\rho N}{2N_2}\cos\theta\,, \qquad (S21)$$

$$\iota_2 = \bar{\iota} - \frac{\rho N}{2N_2}\sin\theta\,. \qquad (S22)$$

Note that $\rho$ and $\theta$ are implicitly restricted by the requirement that $0 \leq \sigma_j \leq 1$ and $0 \leq \iota_j \leq 1$. Under this coordinate transformation, we simply have $h = \rho$, *i.e.* contours of constant heterogeneity are circles centred on $\bar{\sigma}, \bar{\iota}$.

We now turn our attention to *R0*, assuming that $S_j \gg I_j$ and so $N_j \approx S_j$. In our polar coordinates, *R0* becomes

$$R_0 \approx \frac{\kappa}{d + \alpha}(\sigma_1 \iota_1 N_1 + \sigma_2 \iota_2 N_2)$$

$$= \frac{\kappa}{d + \alpha}\left(\left(\bar{\sigma} + \frac{\rho N}{2N_1}\cos\theta\right)\left(\bar{\iota} + \frac{\rho N}{2N_1}\sin\theta\right)N_1 + \left(\bar{\sigma} - \frac{\rho N}{2N_2}\cos\theta\right)\left(\bar{\iota} - \frac{\rho N}{2N_2}\sin\theta\right)N_2\right)$$

$$= \frac{\kappa N}{d+\alpha}\left(\bar{\sigma}\bar{\iota} + \frac{\rho^2}{4}\cos\theta\sin\theta\left(\frac{N}{N_1} + \frac{N}{N_2}\right)\right)$$

$$= \frac{\kappa N}{d+\alpha}\left(\bar{\sigma}\bar{\iota} + \frac{\rho^2}{8}\sin 2\theta\left(\frac{N}{N_1} + \frac{N}{N_2}\right)\right). \qquad (S23)$$

For fixed $\theta$, then, $R_0$ is monotonically increasing or decreasing with $\rho$. Differentiating with respect to $\theta$, we have

$$\frac{\partial R_0}{\partial \theta} = \frac{\kappa N}{d+\alpha}\frac{\rho}{4}\cos 2\theta\left(\frac{N}{N_1} + \frac{N}{N_2}\right), \qquad (S24)$$

which vanishes when $\theta = \frac{\pi}{4}, \frac{3\pi}{4}, \frac{5\pi}{4}, \frac{9\pi}{4}$. Differentiating $R_0$ with respect to $\theta$ again and evaluating at these zeros shows that the first and third of these are maxima and the second and fourth are minima. Thus $R_0$ is maximal for fixed heterogeneity on the positive diagonal $\theta = \frac{\pi}{4}, \frac{5\pi}{4}$, centred on the trait means, in which case we have

$$\sigma_1 - \bar{\sigma} = \iota_1 - \bar{\iota} = \frac{\rho}{2\sqrt{2}N_1}, \qquad (S25)$$

*i.e.* the two traits lie an equal distance in the same direction from the trait means (Figure S1). On the other hand, $R_0$ is minimal for fixed heterogeneity on the negative diagonal $\theta = \frac{3\pi}{4}, \frac{9\pi}{4}$, in which case we have

$$\sigma_1 - \bar{\sigma} = -(\iota_1 - \bar{\iota}) = \frac{\rho}{2\sqrt{2}N_1}, \qquad (S26)$$

*i.e.* the two traits lie an equal distance from, but in opposite directions from, the trait means (Figure S1).

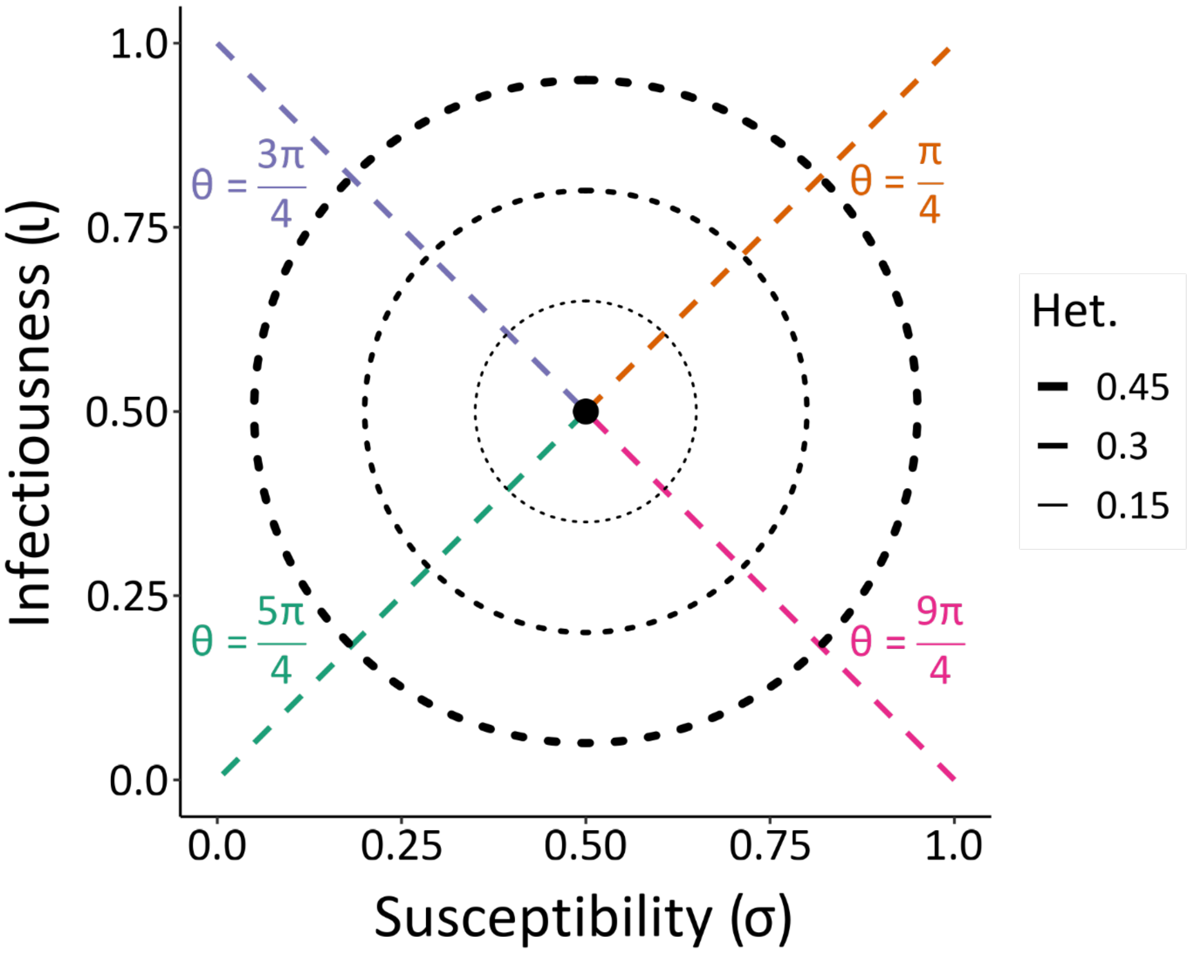


Figure S1. Contours of constant heterogeneity in $\sigma, \iota$ trait space, with two pairs of trait values of equal population abundance. Contours are centred at trait means $\bar{\sigma} = 0.5, \bar{\iota} = 0.5$. The positive diagonal $\theta = \frac{\pi}{4}, \frac{5\pi}{4}$ represents maximal values of $R_0$ on each contour, while the negative diagonal $\theta = \frac{3\pi}{4}, \frac{9\pi}{4}$ represents minimal values of $R_0$.

***Donor dependence***

In this case we assume that $N \approx S_1 + S_2$, as we assume the infected sub-populations are small, so can re-write Equation S16 as

$$R_0 = \frac{\kappa \iota_{\max}}{d + \alpha} \sum_{j=1}^{n_S} \sigma_j S_j \,. \qquad S27$$

We then have

$$\begin{aligned} h &\approx \frac{1}{N}|\bar{\sigma} - \sigma_1|S_1 + \frac{1}{N}|\bar{\sigma} - \sigma_2|S_2 \\ &= \frac{1}{N}|\bar{\sigma} - \sigma_1|S_1 + \frac{1}{N}\left|\bar{\sigma} - \frac{\bar{\sigma}N - \sigma_1 S_1}{S_2}\right| S_2 \\ &= \frac{2}{N}|\bar{\sigma} - \sigma_1|S_1. \qquad (S28) \end{aligned}$$

Hence contours of constant heterogeneity form (approximately) vertical lines in $\sigma_1, \iota_1$ space.

From Equation (12), we have

$$R_0 = \frac{\kappa\bar{\sigma}N}{d+\alpha}\max\left(\iota_1, \frac{1}{I_2}\left(\bar{\iota}(I_1 + I_2) - \iota_1 I_1\right)\right), \qquad (S29)$$

where we have used Equation (S6) to write $\iota_2$ in terms of $\bar{\iota}$. Now, $\max\left(\iota_1, \frac{1}{I_2}\left(\bar{\iota}(I_1 + I_2) - \iota_1 I_1\right)\right) = \iota_1$ only if $\iota_1 \geq \bar{\iota}$, so in this case $R_0$ increases monotonically as $\iota_1$ moves away from $\bar{\iota}$ in either direction, as seen in Figure 2. The case shown in Figure 2 has $\bar{\sigma}$ fixed at 0.5, so there is no variation in the $\sigma_1$ direction.

### 3. Bipartite heterogeneity results

#### ***RD host abundance***

Enlarging Figure 3 (main text) shows that total abundance is at the lowest values when the minimum population-level susceptibility value ($\sigma_{min}$) is at its highest, regardless of the amount of heterogeneity (Figure S2B).

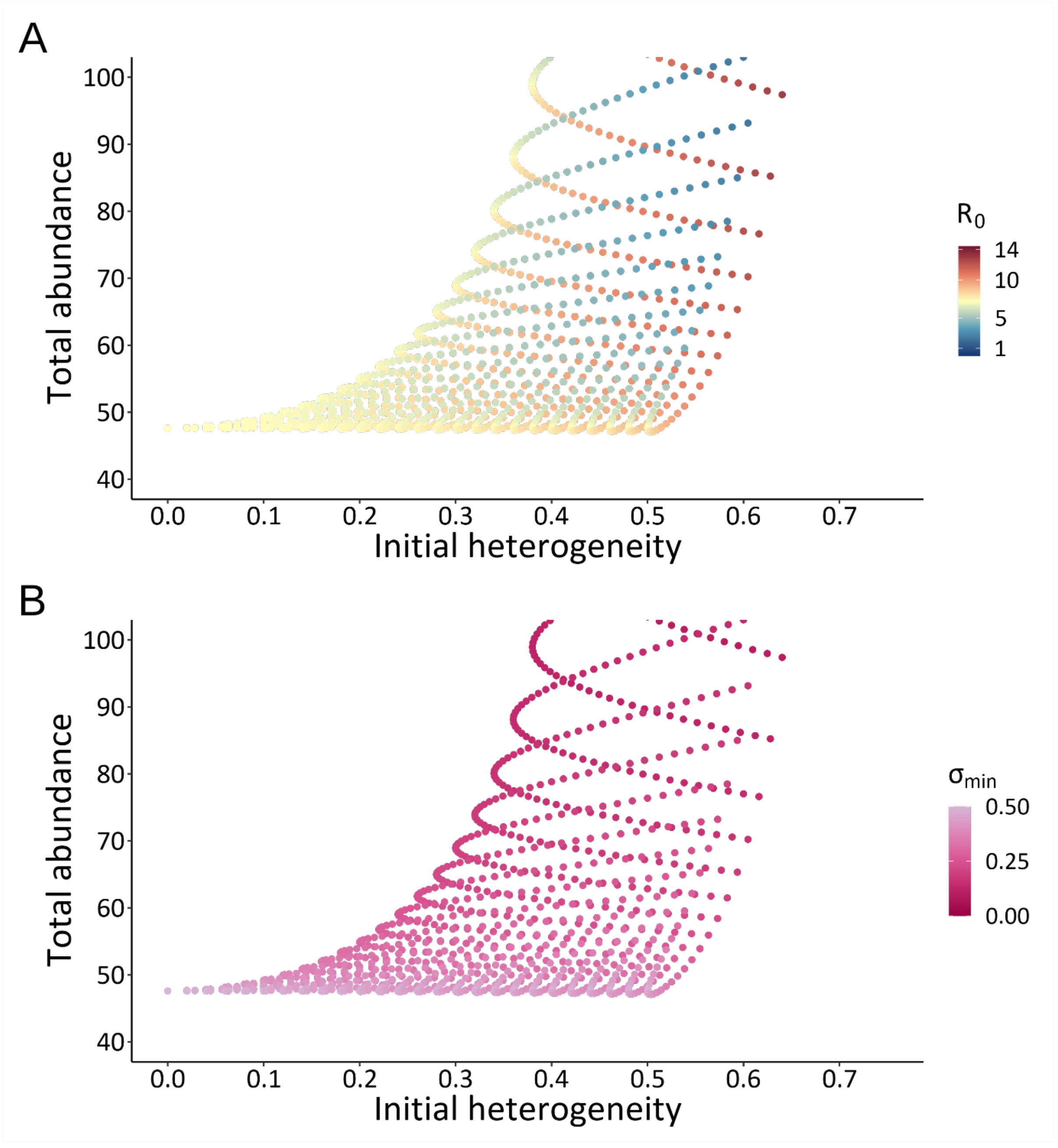


Figure S2. Effect of initial heterogeneity on equilibrium total host abundance for RD, enlarged to focus on total abundances below 100. In (A) the colour bar shows $R_0$ values. In (B) the colour bar shows the minimum population-level susceptibility value ($\sigma_{min}$).

***Change in heterogeneity***

To understand how epidemic progress affects population-level heterogeneity we compared initial heterogeneity and its value at equilibrium. For both the RD and DD scenarios there is generally very little change in population-level heterogeneity throughout the epidemic (Figure S3).

Counterintuitively, we did not find a noticeable divergence between the RD and DD scenarios in how population-level heterogeneity changed during an epidemic. In the DD scenario we expected to see a loss of heterogeneity over time because the infected sub-population with the highest infectiousness value in the population becomes dominant as the epidemic progresses, ultimately excluding less infectious donors. However, heterogeneity is calculated by taking the mean abundance-weighted distance of the sub-populations to the centroid. Thus, despite the DD scenario losing infected sub-populations at equilibrium (and leading to maximally infectious hosts over time), the weighting of the heterogeneity score with the generally larger susceptible sub-populations ensures that there is no considerable loss in heterogeneity at equilibrium.

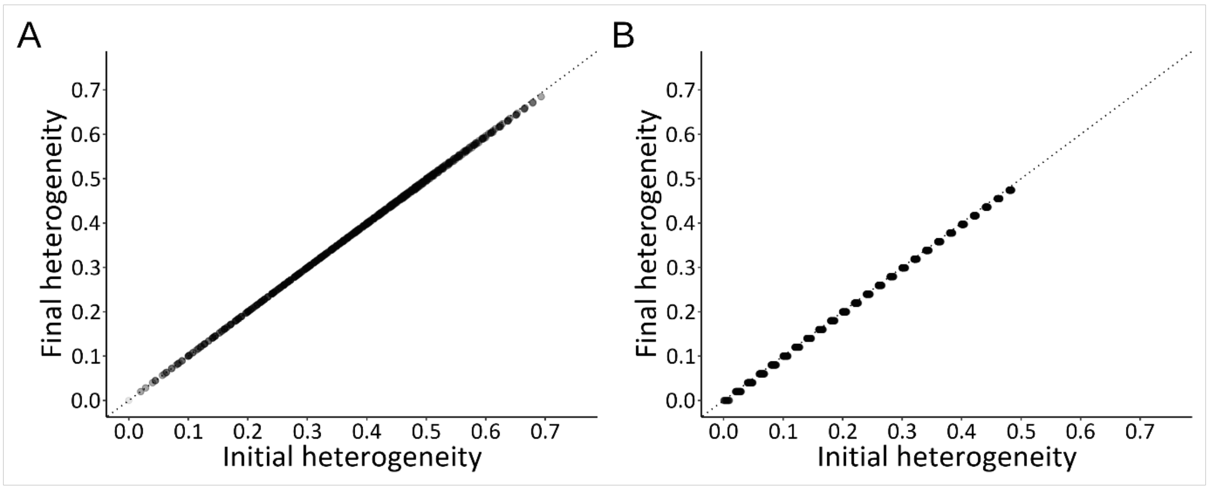


Figure S3. Change in heterogeneity from the initial sub-population values to their equilibrium values (the dotted line shows $y = x$ in both panels). (A) RD scenario, (B) DD scenario. Each point represents one simulation; points are light grey, such that darker points represent multiple, overlapping points.

### 4. Tripartite heterogeneity results

The results for the analyses of the tripartite heterogeneity contexts 2 and 3 are presented here. For each context we generated a range of unique combinations of $\sigma$ and $\iota$ trait values (n = 8,089 tripartite isometric heterogeneity, n = 109,561 tripartite non-isometric heterogeneity). The results are generally consistent with those from the bipartite heterogeneity analyses, particularly in the case of the tripartite non-isometric heterogeneity results, and are described in more detail below.

#### Heterogeneity and $R_0$

##### *Recipient dependence*

When three sub-populations are equally spaced in $\sigma/\iota$ trait space (*i.e.* tripartite isometric context, with a centroid fixed at ($\sigma = 0.5, \iota = 0.5$)), the covariation between susceptibility ($\sigma$) and infectiousness ($\iota$) is 0, regardless of the amount of heterogeneity present. Because the covariation of $\sigma$ and $\iota$ is 0, $R_0$ is the same in the homogeneous instance as all heterogeneous simulations (Figure S4).

Increasing heterogeneity in the tripartite non-isometric heterogeneity context leads to divergent results for $R_0$ (Figure S4C), as was seen in the bipartite heterogeneity context. When there is positive covariation between susceptibility ($\sigma$) and infectiousness ($\iota$), $R_0$ values increase; negative covariation reduces $R_0$ values (Figure S4C). Again, as for the bipartite heterogeneity case, the shape of the point cloud is constrained by how sub-populations can be positioned in $\sigma, \iota$ trait space.

##### *Donor dependence*

With equally spaced sub-populations (tripartite isometric heterogeneity), increasing heterogeneity in infectiousness increases $R_0$ values (Figure S4B). This is because $R_0$ values are determined by the maximum infectiousness ($\iota$) value. These results are broadly similar to the bipartite heterogeneity case.

Assuming tripartite non-isometric heterogeneity, the maximum infectiousness ($\iota$) value in each simulation determines the $R_0$ value, leading to a general increase in $R_0$ with increasing heterogeneity (Figure S4D). Though there is no change in $R_0$ if the maximum infectiousness ($\iota$) value does not change as heterogeneity increases (*i.e.* only an increase in heterogeneity in susceptibility ($\sigma$)). These findings are once again consistent with both the bipartite heterogeneity and the tripartite isometric heterogeneity contexts.

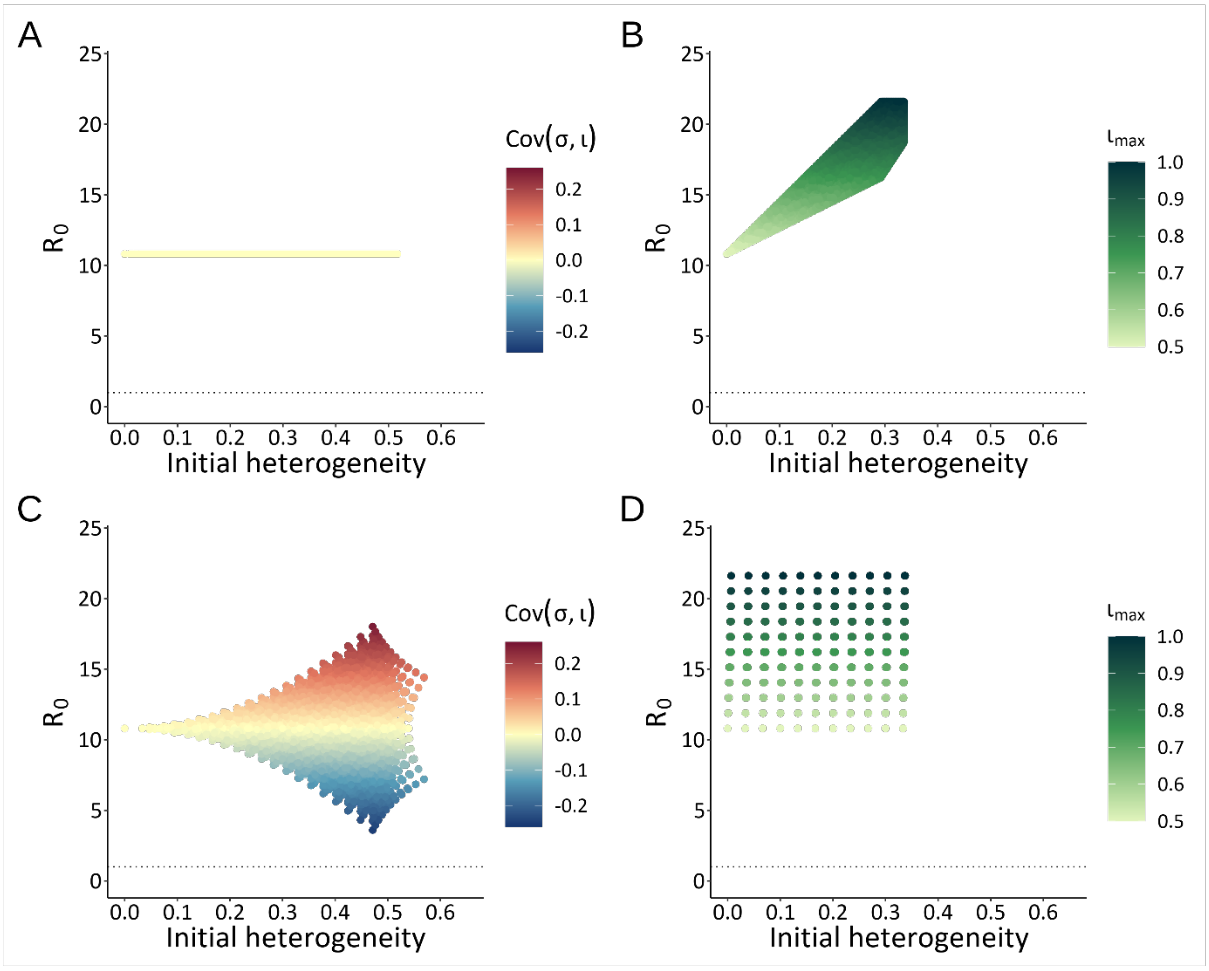


Figure S4. The effect of initial heterogeneity on $R_0$. A) Results from the RD tripartite isometric heterogeneity context, where each point represents a unique simulation. The colour bar shows the covariance of susceptibility ($\sigma$) and infectiousness ($\iota$) in each simulation. B) Results from DD tripartite isometric heterogeneity, where each point represents a unique simulation. The colour bar shows the maximum infectiousness ($\iota$) value in each simulation. C) Results from RD tripartite non-isometric heterogeneity, where each point represents a unique simulation. The colour bar shows the covariance of susceptibility ($\sigma$) and infectiousness ($\iota$) in each simulation. D) Results from DD tripartite non-isometric heterogeneity, where each point represents a unique simulation. The colour bar shows the maximum infectiousness ($\iota$) value in each simulation. In all panels the dotted line shows where $R_0 = 1$.

**Host abundance**

In both the RD and DD tripartite isometric heterogeneity context, equilibrium host abundance increases with greater initial heterogeneity (Figure S5A & Figure S5B). Some total abundances increase by a much larger magnitude as heterogeneity values increase, while most increase by a smaller magnitude. While the DD tripartite isometric heterogeneity context (Figure S5B) is broadly similar to that of its RD counterpart, the lowest abundances are found at median levels of heterogeneity.

Similarly, the RD and DD tripartite non-isometric heterogeneity contexts show greater total host abundance with increasing heterogeneity (Figure S5C & Figure S5D). As with the bipartite heterogeneity abundance results (main text, Figure 4), simulations with higher $R_0$ values have lower total abundances, and simulations are grouped.

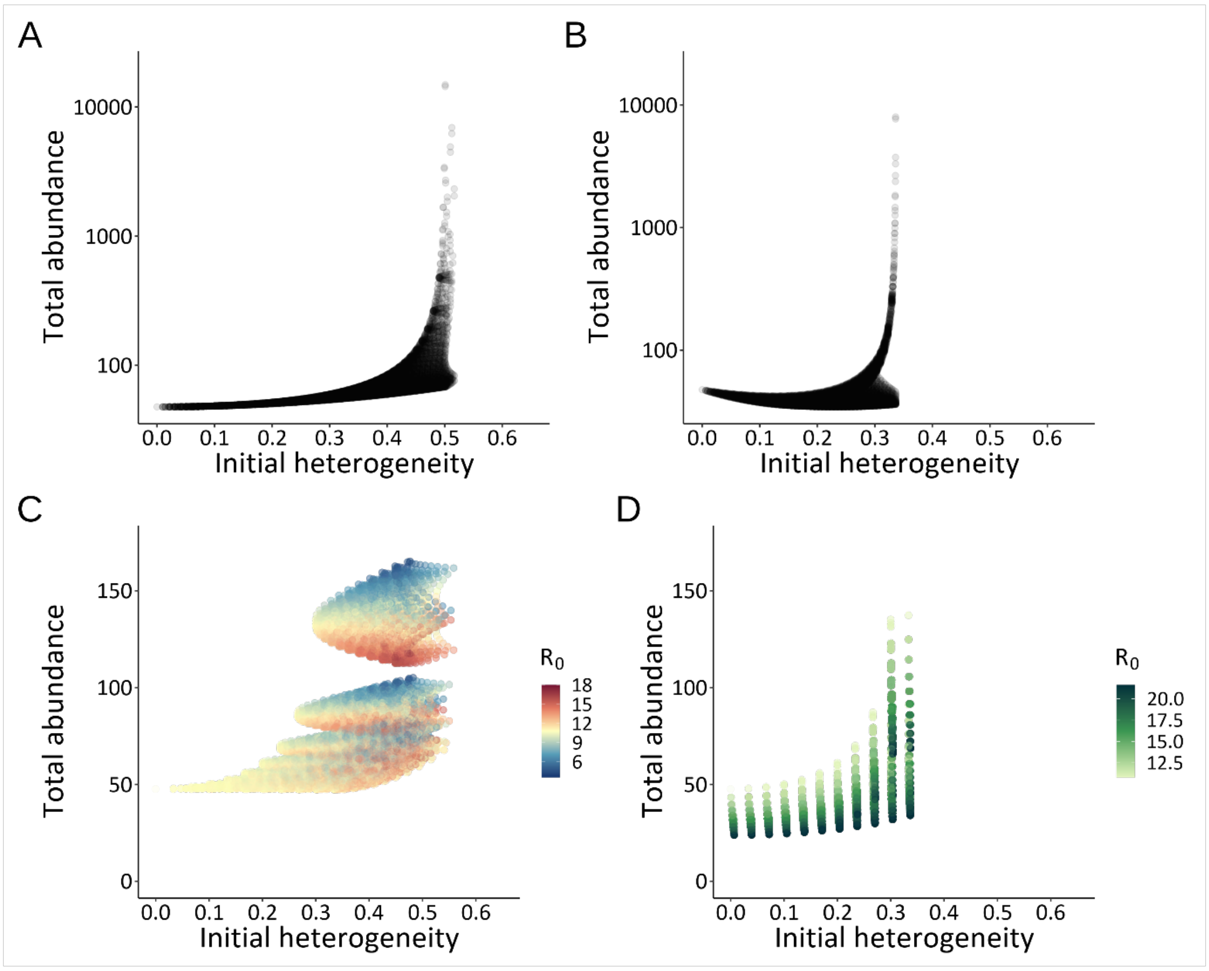


Figure S5. Effect of initial heterogeneity on total abundance of hosts at equilibrium. A) RD tripartite isometric heterogeneity context. B) DD tripartite isometric heterogeneity context. C) RD tripartite non-isometric heterogeneity context. D) DD tripartite non-isometric heterogeneity context.

**Change in heterogeneity**

Both the RD and DD tripartite non-isometric heterogeneity contexts broadly show the same pattern as for the bipartite heterogeneity context (compare Figure S6 with Figure S3). Although there is slightly more deviation from the $y = x$ line in both the RD tripartite non-isometric heterogeneity and the DD tripartite isometric heterogeneity context when compared to the RD tripartite isometric heterogeneity or the DD tripartite non-isometric heterogeneity cases, the general pattern shows that heterogeneity tends to be maintained at equilibrium (Figure S6).

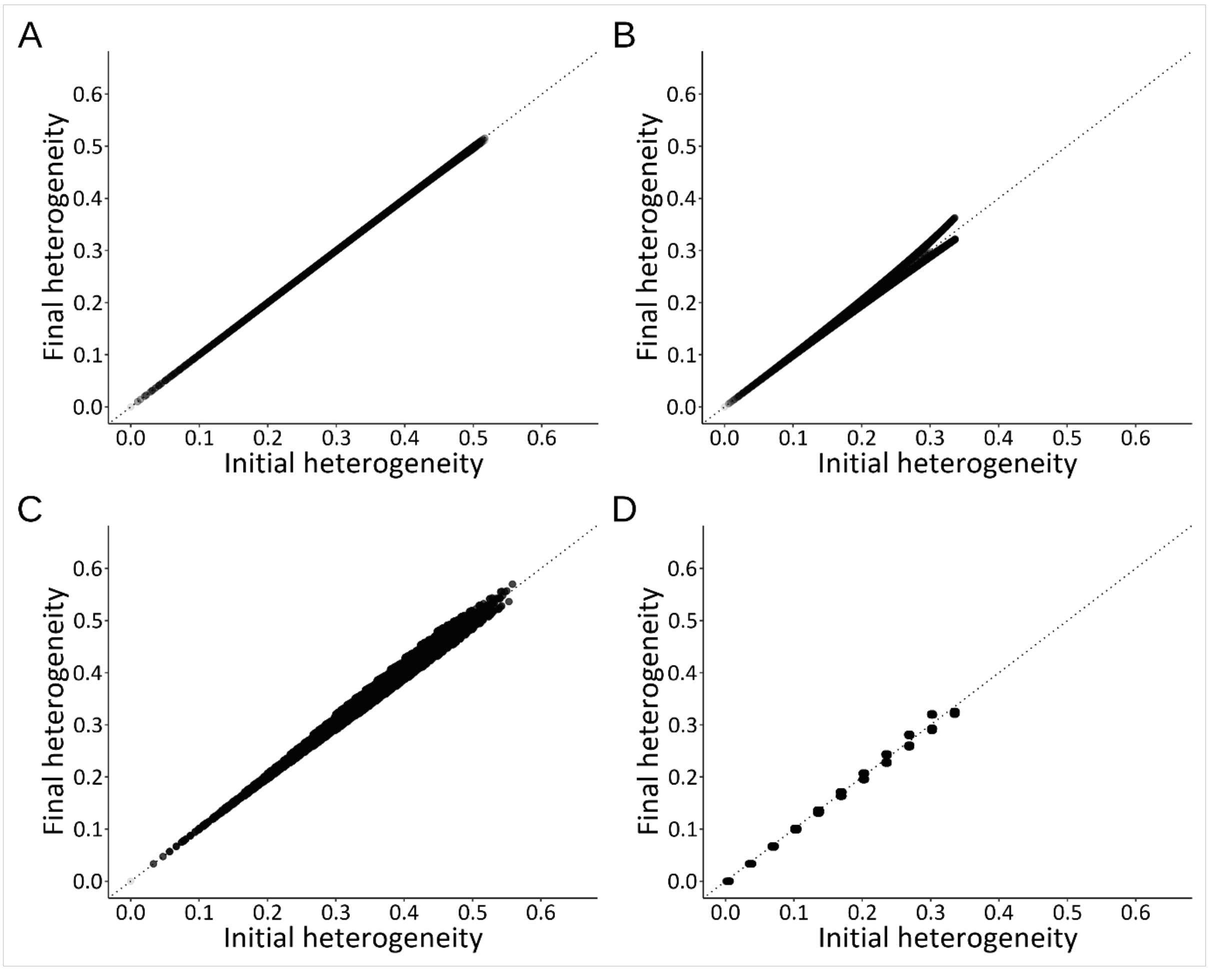


Figure S6. Change in heterogeneity from the initial sub-population values to their equilibrium values. In each panel the dotted line shows $y = x$, each point represents at least one simulation, and darker points are multiple overlaid simulations. A) RD tripartite isometric heterogeneity context. B) DD tripartite isometric heterogeneity context. C) RD tripartite non-isometric heterogeneity context. D) DD tripartite non-isometric heterogeneity context.

### References

Diekmann, O., Heesterbeek, J. & Roberts, M.G. (2010). The construction of next-generation matrices for compartmental epidemic models. *Journal of the Royal Society Interface*, 7, 873-885.

Laliberté, E. & Legendre, P. (2010). A distance-based framework for measuring functional diversity from multiple traits. *Ecology*, 91, 299-305.
